# Longitudinal increase in sleep problems is related to amyloid deposition in cortical regions with high *HOMER1* gene expression

**DOI:** 10.1101/335612

**Authors:** Anders M Fjell, Donatas Sederevicius, Markus H Sneve, Ann-Marie Glasø de Lange, Anne Cecilie Sjøli Bråthen, Kristine B Walhovd, for The Alzheimer’s Disease Neuroimaging Initiative

**Affiliations:** Center for Lifespan Changes in Brain and Cognition, Department of Psychology, University of Oslo, Oslo, Norway; Department of radiology and nuclear medicine, Oslo University Hospital, Oslo, Norway

## Abstract

Older adults who report more sleep problems tend to have elevated levels of the Alzheimer’s disease (AD) biomarker β-amyloid (Aβ), but the mechanisms responsible for this relationship are largely unknown. Molecular markers of sleep problems are now emerging from rodent research, yielding opportunities to generate hypotheses about the causes of the sleep-Aβ relationship. A major molecular marker of sleep deprivation is Homer1a, a neural protein coded by the *HOMER1* gene, involved in control of sleep homeostasis and also implied in Aβ accumulation. Here, in a sample of 109 cognitively healthy middle-aged and older adults, we tested whether the relationship between cortical Aβ accumulation and self-reported sleep quality, as well as changes in sleep quality over three years, was stronger in cortical regions with high *HOMER1* mRNA expression levels. Aβ correlated with poorer sleep quality cross-sectionally and longitudinally. This relationship was stronger in the younger (50-67 years) than the older (68-81 years) participants. Effects were mainly found in regions with high expression of *HOMER1*, suggesting a possible molecular pathway between sleep problems and Aβ accumulation. The anatomical distribution of the sleep-Aβ relationships followed closely the Aβ accumulation pattern in 69 patients with mild cognitive impairment (MCI) or AD. Thus, the results indicate that the relationship between sleep problems and Aβ-accumulation may involve Homer1 activity in the cortical regions that harbor Aβ in AD. Analysis of cortical gene expression patterns represent a promising avenue to unveil molecular mechanisms behind the relationship between sleep problems and AD risk.

## Introduction

Sleep is critical for normal brain function, and disturbed sleep is associated with many prevalent neurological conditions. For instance, sleep disturbance is an early symptom of Alzheimer’s disease (AD) (1-3), and a bidirectional relationship is suggested between sleep quality and β-amyloid (Aβ) accumulation (4). Sleep quality is also reduced in normal aging (5), and understanding the relationship between sleep and Aβ-accumulation in cognitively healthy older adults could aid us in understanding the role of sleep in early stages of neurodegeneration. Although a relationship between sleep and Aβ is demonstrated in experimental rodent studies (6), correlations found in humans are usually moderate or week (7-10).

Here we approach the question of how sleep and amyloid accumulation are related from three different angles. First, accumulation of Aβ is assumed to be an early event in AD, possibly plateauing even before the stage of diagnosis (11-14). Relationships between Aβ and sleep problems may therefore be stronger among “young-older” than “old-older” adults. Second, since sleep problems usually tend to be relatively stable over time (15), increase in sleep problems over time may be a more alarming symptom reflecting initiation of brain pathology. How longitudinal worsening of sleep problems relates to Aβ accumulation is not known, however. Finally, sleep loss has a profound effect on expression of selective genes. Since gene expression varies across the cerebral cortex, testing how the regional distribution of sleep-Aβ relations maps onto expression of genes sensitive to sleep could be a useful approach to indirectly target molecular mechanisms for Aβ-sleep relationships in humans. The best known molecular marker of sleep need is the Homer1a (16, 17), widely expressed in the human cortex, especially in the frontal lobe (18). Homer1 is a neuronal protein coded by the *HOMER1* gene, classified as an immediate early gene. Its expression is broadly upregulated in the brains of sleep deprived rodents (19, 20), induced by sustained neuronal activity when awake (18, 21, 22). Studies of transgenic mice show consistent activation of Homer1a when awake, indicating a role for sleep in intracellular calcium homeostasis for protecting and recovering from glutamate-induced neuronal hyperactivity imposed by wakefulness (16). A recent study suggested that Homer1a serves as a molecular integrator of arousal and sleep via the wake-promoting neuromodulator noradrenaline and the sleep-promoting neuroregulator adenosine, thereby scaling down excitatory synapses during sleep (23). Importantly, because Homer 1a is activity-dependent, it has been suggested to regulate Aβ toxicity at the early stage of AD (24), and reduced Homer 1a mRNA expression has been found in the amyloid precursor protein and presenilin-1 (APP+PS1) transgenic mice (25). In this study, normal expression was found in regions that did not accumulate Aβ, implying a role for Homer1a in Aβ processing. An experimental study further showed that activity-dependent expression of Homer1a counteracted suppression of large-conductance Ca^2+^-activated K^+^ (BK - Big Potassium) channels, demonstrated by injections of Aβ proteins into rat and mouse neocortical pyramidal cells (26). Aβ is assumed to result from synaptic activity, which can explain why Aβ accumulations in humans are preferably found in multi-modal brain regions that show continuous levels of high neural activity and plasticity across the lifespan (27). Thus, a first step in exploring the role of *HOMER1* for Aβ accumulation in humans is to test how its mRNA expression levels in the cerebral cortex maps onto Aβ accumulation in AD and the regional distribution of sleep-Aβ relationships in aging. We would then expect that *HOMER1* in Aβ negative individuals is upregulated in the same cortical regions where Aβ accumulation due to sleep problems or AD is found.

In the present study, 109 cognitively normal older adults (mean age 66.7 years, range 49.2-80.9 years, see Table 1) completed the Pittsburgh Sleep Quality Index (PSQI) (28) and underwent Flutemetamol (^18^F) positron emission tomography (PET) for quantification of cortical Aβ accumulation. 62 of the participants were followed for three years with longitudinal PSQI. The participants underwent extensive neuropsychological and somatic testing and assessment of depressive symptoms (29). We hypothesized that sleep problems, and in particular increases in sleep problems over time, would be more strongly associated with Aβ accumulation in the younger (< 68 years) than the older (> 68 years) part of the sample. We further hypothesized that this relationship would be most evident in AD-vulnerable regions, and in regions with high levels of *HOMER1* mRNA expression.

**Table 1.** Sample descriptives. Beck: Available for 82. Registered 3 years before baseline in the longitudinal sample for 61 BMI: Available for 60 at baseline in the longitudinal subsample Rey available for 87 in the full sample, and 61 in the longitudinal sample. Follow up was at a mean of 3 years after baseline.

|  | Full sample (n = 109) | Longitudinal PSQI (n = 62) |  |
| --- | --- | --- | --- |
|  |  | Baseline | Follow-up |
| Age | 66.7 (49.2-80.9) | 62.1 (8.3) | 66.4 (7.9) |
| Sex (female/ male) | 64/ 45 | 37/25 | 27/25 |
| MMSE | 28.8 (25-30) | 29.0 (27-30) | 28.5 (25-30) |
| PSQI Global | 5.0 (2.77) | 4.5 (2.73) | 4.5 (2.77) |
| BMI | 25.8 (3.96) | 26.0 (4.21) | 25.5 (3.80) |
| Beck (BDI) | 0-18 (5.1) | 5.11 (3.82) | 4.48 (4.30) |
| WASI vocabulary (t-score) | 59.9 (6.6) | 59.4 (7.1) | 59.9 (5.9) |
| WASI matrices (t-score) | 63.0 (8.3) | 62.4 (7.9) | 63.9 (7.9) |
| Rey Complex Figure (memory) | 18.0 (6.3) | 17.9 (5.4) | 17.7 (6.2) |

## Results

### Global amyloid levels – cross sectional analyses

PET scans were partial voluming corrected and mapped to the reconstructed cortical surface of each participant (30). The cortex was parcellated into 34 regions in each hemisphere (31), and standardized uptake values (SUV) for each region were calculated as the ratio between the region value and the mean value of the cerebellum gray matter. Global cortical Aβ accumulation was calculated by use of principal component analysis of the 68 regions, yielding one standardized measure for each participant accounting for 68.1% of the variance in Aβ (see Supplemental Information). The PSQI global score was used as measure of sleep quality.

Global cortical Aβ (r =.19, p <.05) correlated with age, while PSQI (r =.10, p =.29) did not, probably because only middle-aged and older adults were included. PSQI was entered as the dependent variable in a multiple regression analysis, with Aβ, age and sex as predictors. Aβ and age were not related to PSQI in this model (p >.50), while sex showed a trend towards more sleeping problems for females (β = -.17, p =.092). The Aβ × age interaction term was added, yielding a significant contribution to explain PSQI (β = -.22, p <.05). Post hoc partial correlation analyses, controlling for age, showed that this interaction was due to a positive relationship (r =.41, p <.005) between sleep problems, i.e. high PSQI score, and Aβ in the youngest (age < 68 years) part of the sample and no significant relationship (r = -.23, p =.084) in the oldest (age > 68 years) (Figure 1). As depression symptoms are associated with more sleep problems, we re-ran the model with score on the Beck Depression Inventory (BDI) (29, 32) as an additional covariate. Aβ × age was still significant (β = -.29, p <.05), as was BDI score (β =.40, p <.001), meaning that more sleep problems were associated with more depressive symptoms. Further, overweight and cardiovascular health can affect sleep quality (33). The Aβ × age interaction survived including body mass index (BMI) as an additional covariate (β = -.22, p <.05), and BMI did not contribute significantly. In addition to BMI, we also added a range of measures related to body composition obtained from bioelectrical impedance examinations (34) (muscle mass, body fat percentage, waist circumference, waist/ hip ratio, visceral fat area), as well as blood pressure (systolic and diastolic). None of these explained any unique variance in PSQI score, while the Aβ × age interaction was still significant (β = -.29, p <.05).

**Figure 1.**
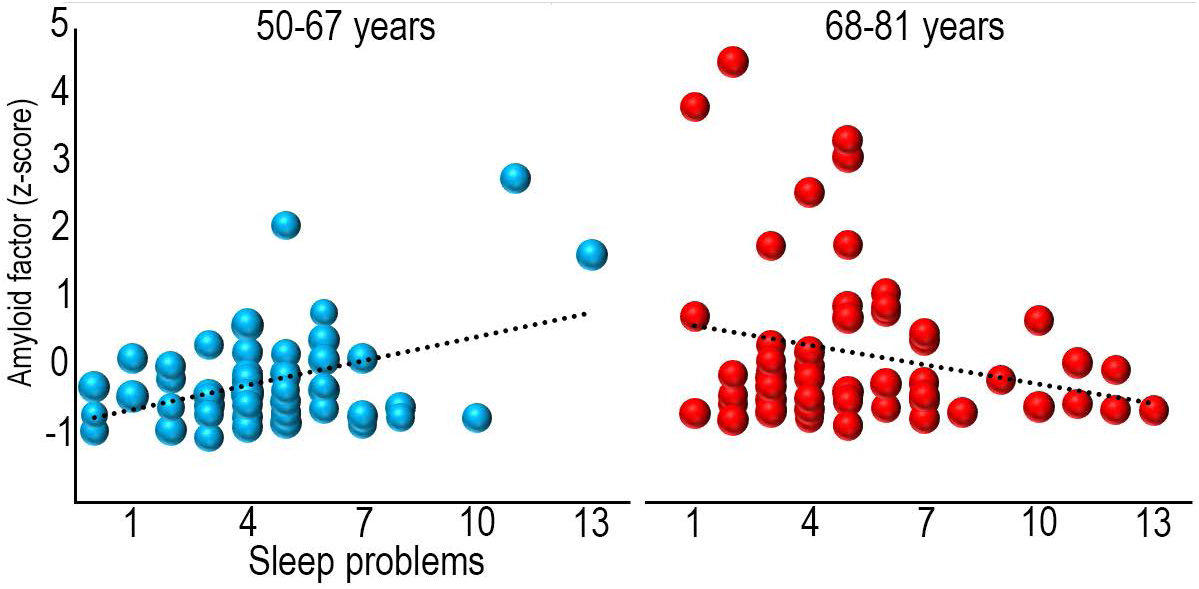
Sleep problems and global Aβ levels. Bubble plots of the relationship between sleep problems (PSQI global score) and global cortical Aβ levels (amyloid factor expressed in Z-scores) within each age group. The size of the bubbles within each age group are scaled with the age of the participants.

### Regional amyloid levels – cross-sectional analyses

Next, we tested for regional specificity in the PSQI amyloid relationships. General linear models (GLM) were run with Aβ level at each cortical vertex as dependent variable, and age group, PSQI score and age as predictor variables. The results showed spatially extended age group × PSQI interactions (Figure 2), covering the frontal lobes, lateral temporal lobes, inferior parietal cortex as well as the medial parietal cortex. The relationships were significantly stronger in participants below 68 years (Figure 3, see SI for scatterplots).

**Figure 2.**
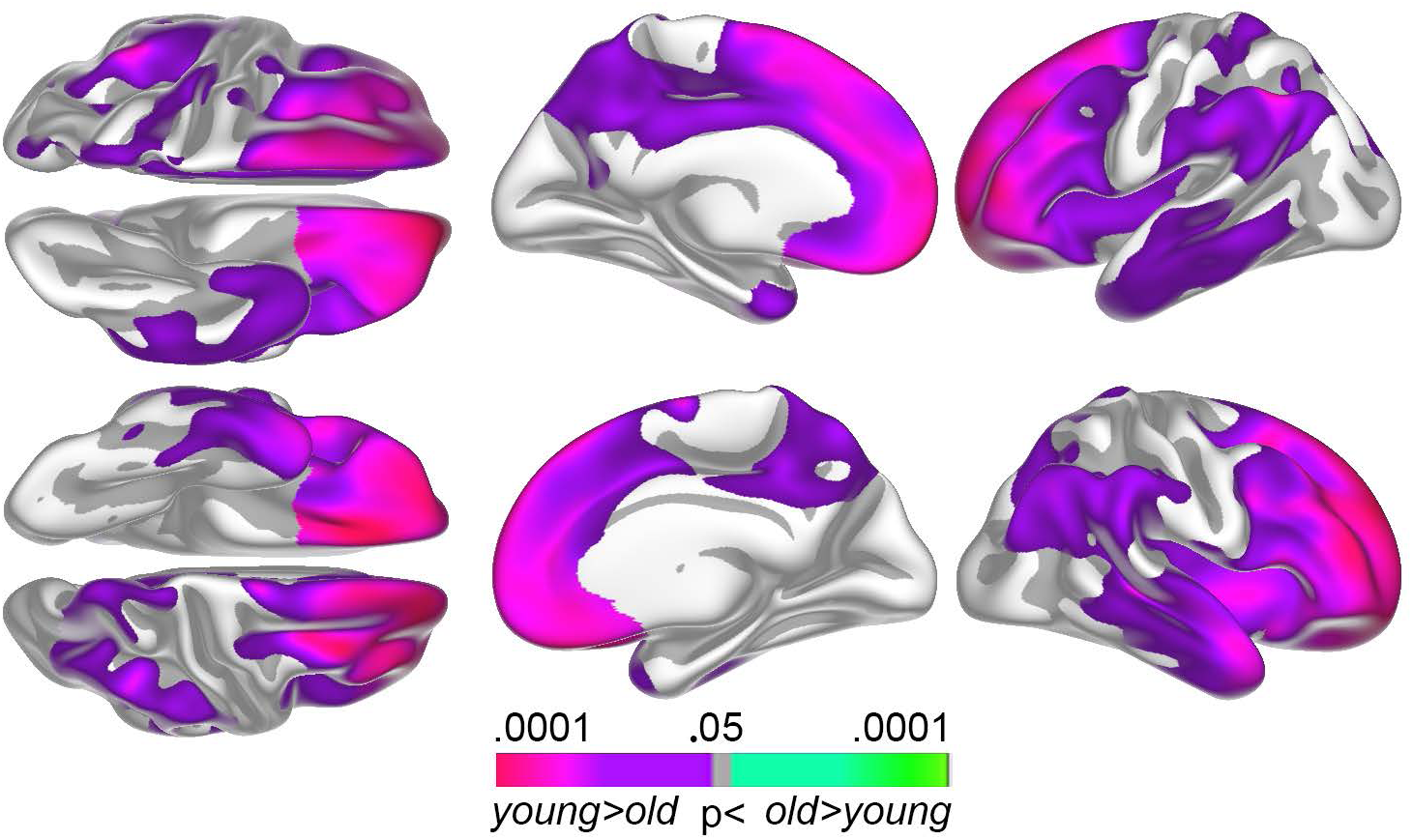
Regional Aβ levels and sleep problems. The surface plots show regions where sleep problems and Aβ accumulation are significantly stronger correlated in the younger (50-67 years) compared to the older (68-81 years) age group.

**Figure 3.**
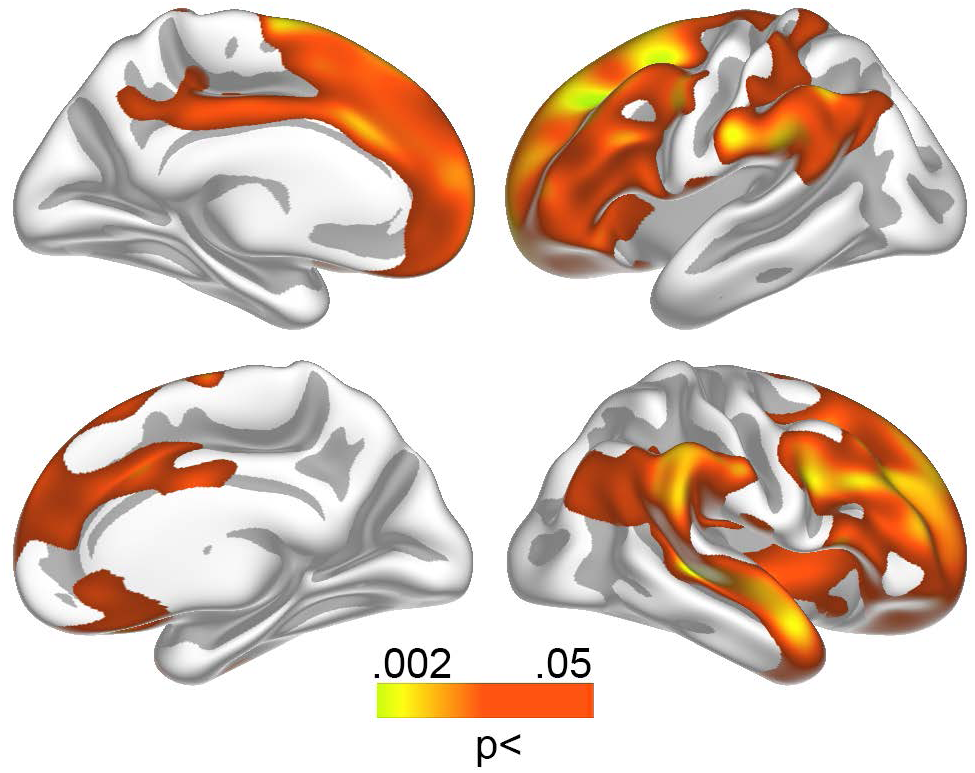
Relationships between Aβ levels and sleep problems in the youngest group. Relationship between sleep problems and Aβ accumulation in the youngest participants (50-67 years).

### Global and regional amyloid levels – longitudinal analyses

PSQI, memory and depression scores were available for a subsample from three years prior to the present investigation (n = 62, mean interval = 3.0 years). PSQI did not change significantly over this time (mean change = -4.0%, SD = 29%, t = 1.16, n.s.), and PSQI scores between time points were highly correlated (r =.81, p < 10^-14^). A regression model with PSQI change as dependent variable, and age group, global Aβ accumulation and the age × Aβ interaction as independent variables, showed that the interaction term was significant (β =.23, p =.05). Worsening of sleep problems over time correlated with higher Aβ levels in the youngest (r =.37, p <.05, n = 35) but not the oldest (r = -.12, n.s, n = 27) age group. The relationship in the young group was upheld if baseline PSQI score was added as a covariate (r =.40, p <.05).

### Alzheimer’s Disease-dependent Aβ accumulation

We tested whether sleep-related Aβ accumulated in the AD-vulnerable regions. If this was the case, it could mean that the observed correlations between sleep problems and Aβ are relevant for understanding early stages of AD pathophysiology. However, if the overlap between the Aβ – PSQI relationships and AD-related Aβ-accumulation was low, this would indicate that the relationships more likely represent processes independent of AD. Thus, we compared cortical Aβ accumulation in 20 cognitively healthy older adults (age 71.3-86.2 years) with 69 patients with MCI/ AD (n = 44/ 25, age 55.3-88.2 years) randomly drawn from the Alzheimer’s Disease Neuroimaging Initiative (ADNI; adni-info.org) (Figure 4). As expected, the patients harbored significantly more cortical Aβ than the controls. The regional variability in the amount of difference between patients and controls was substantial, being especially pronounced in the superior frontal gyrus and around the central sulcus. Aβ did not differ between patients and controls in large parts of the occipital and medial temporal cortex. A direct comparison of the anatomical distribution of Aβ differences between controls and patients (gamma values, i.e. simple effect size) with the distribution of the sleep-related Aβ accumulation revealed a close overlap (Spearman’s Rho =.81, p < 10^-8^), clearly showing that sleep is related to Aβ in the AD-sensitive regions.

**Figure 4.**
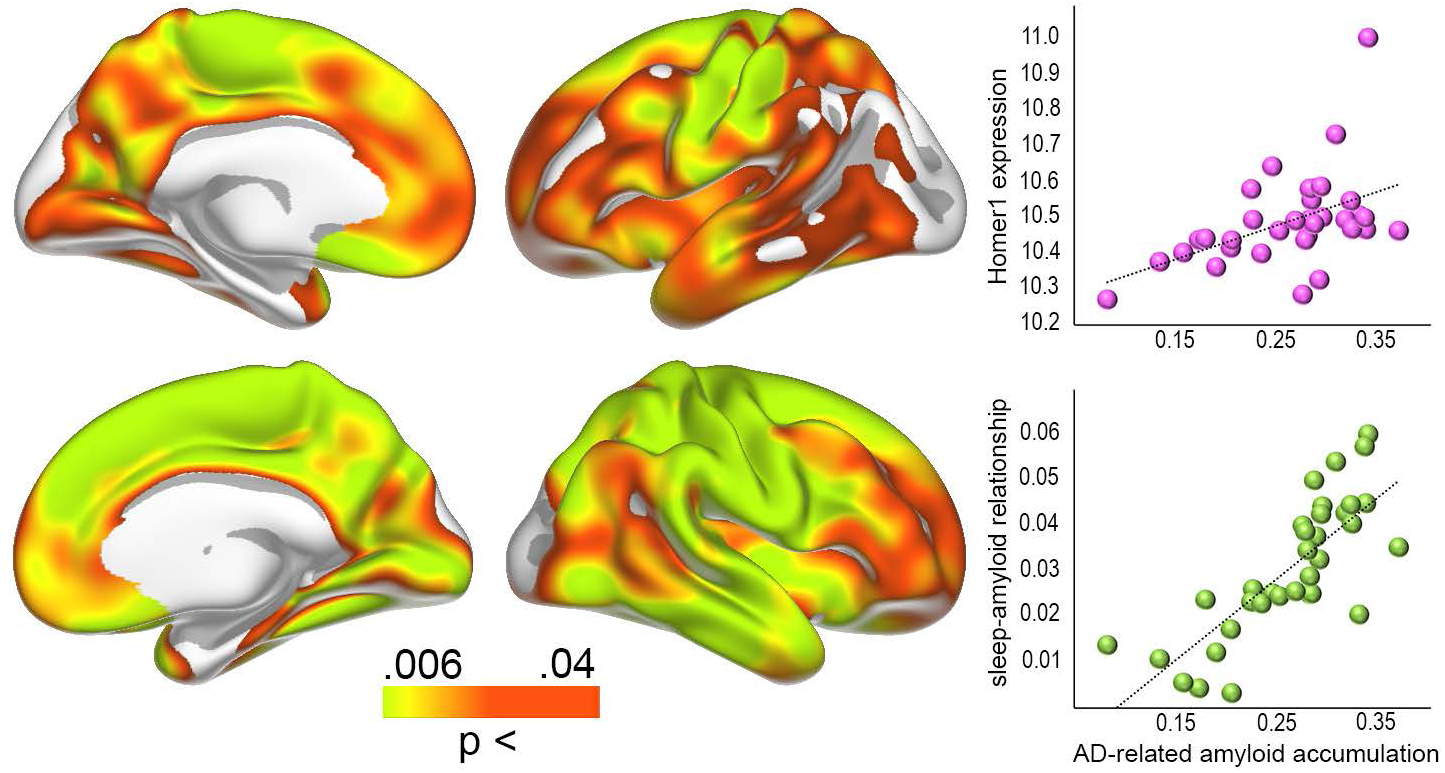
Aβ levels in AD and gene expression. Left panel: regions with significantly higher levels of Aβ in MCI/ AD patients compared to cognitively normal controls. Right panel: Bubble plot of the relationship between the patients vs. controls differences in Aβ accumulation across 34 cortical regions (gamma values) and regional *HOMER1* expression levels (top) and strength of the sleep-Aβ relationship in the youngest group (gamma values) vs. the patients-controls differences (bottom).

### Gene expression – HOMER1

Anatomical distribution of the mRNA expression levels of *HOMER1* from the Allen Brain Atlas (35) is shown in Figure 5. The analyses were restricted to the left hemisphere because the brain atlas consists of more donors for the left (n = 6) than the right (n = 2) hemisphere. For each of the 34 cortical regions, we extracted normalized gene expression for *HOMER1* (36), see SI. Median *HOMER1* expression across donors was then correlated with the effect size from the cross sectional Aβ – PSQI analyses in the youngest group (shown in Figure 3). A significant correlation was found (Spearman’s Rho =.51, p =.0022, see Figure 5). As an additional test, we ran the analysis for all 20736 genes in the atlas and compared the observed *HOMER1* expression correlation to the distribution of all possible gene expression - Aβ – PSQI correlations. The observed correlation of.51 for *HOMER1* was well above 97.5% of all positive correlations (critical value of Spearman’s Rho =.49), demonstrating that *HOMER1* is among the top 5% correlated genes, similar to a two-tailed p <.05. Control analyses showed that all donors showed similar positive correlations when investigated individually (see SI).

**Figure 5.**
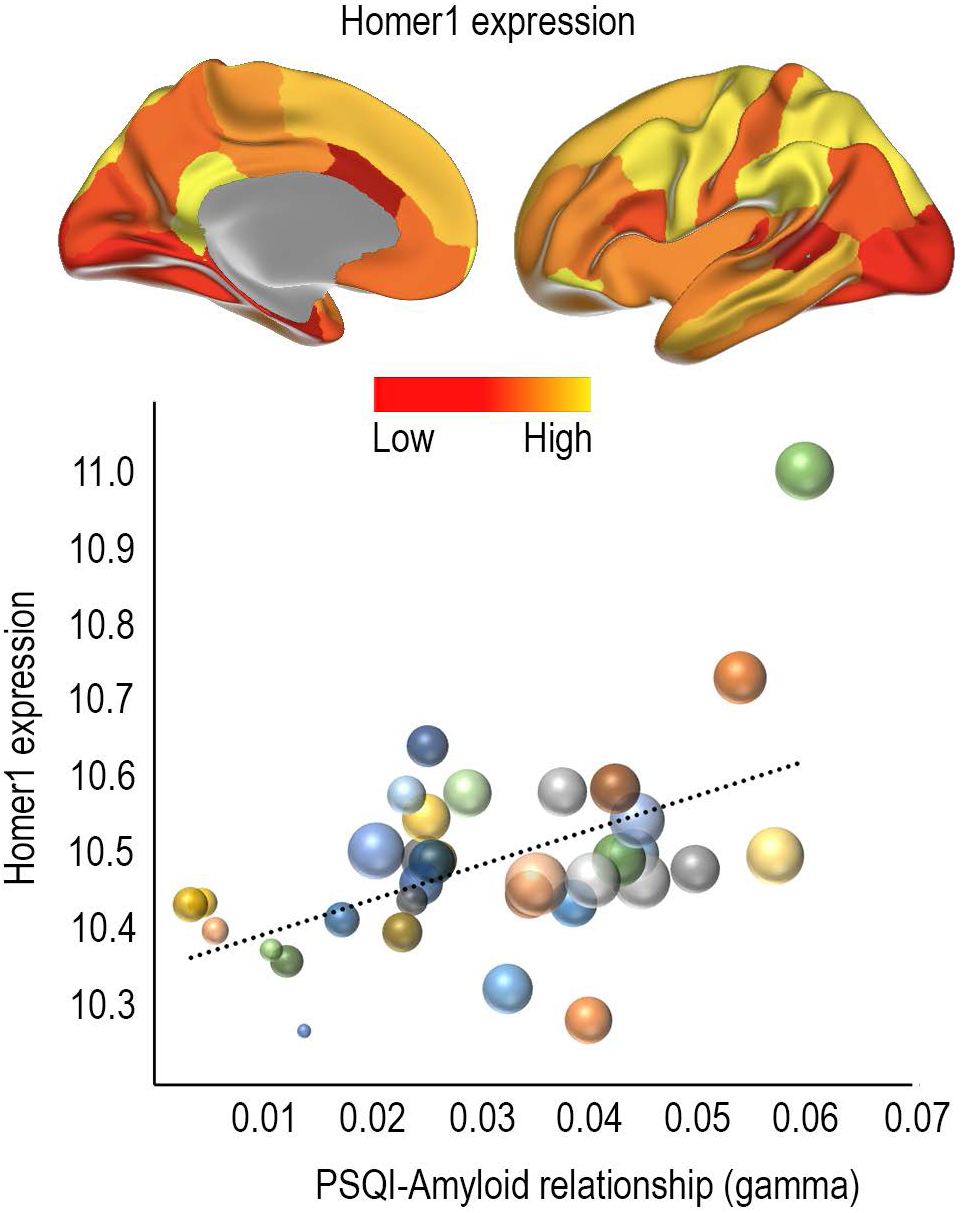
Relationship between sleep problems related Aβ accumulation and *HOMER1* expression. Top panel: Regional expression of *HOMER1* in 34 cortical regions in the left hemisphere. Bottom panel: Bubble plot of the relationship between regional *HOMER1* expression levels and strength of the sleep-Aβ relationship in the youngest group (gamma values). The bubbles are scaled by the group difference in Aβ accumulation between controls and MCI/ AD patients. The clustering of large bubbles to the right and to the top of the plot illustrates that regions with high levels of *HOMER1* expression and sleep-related Aβ accumulation also show more Aβ accumulation in AD patients.

The results indicate that *HOMER1* expression and sleep may be relevant players in very early Aβ accumulation in humans. A role for Homer1 in Aβ accumulation is also suggested by rodent work showing reduced expression of Homer 1a mRNA in regions accumulating Aβ in APP+PS1 transgenic mice and normal expression in regions without Aβ accumulation(25). Based on such findings, if Homer1 is a key player in the relationship between sleep problems and AD pathology, we would expect strong expression of *HOMER1* in cognitively healthy Aβ negative individuals in the regions where AD pathology accumulates. As hypothesized, *HOMER1* expression was positively related to AD-related Aβ accumulation (Spearman’s Rho =.55, p <.001, > 99^th^ percentile compared to all genes, see Figure 4).

### Control analyses - Sleep, memory and depression

As sleep is related to depression and memory function, we ran control analyses testing the relationship between PSQI scores, global Aβ, memory function (The Rey-Osterreith Complex Figure Text, a visuo-constructive recall test (37)) and symptoms of depression (29). Controlling for age, baseline PSQI score correlated negatively with recall (r = -.34, p <.01, df = 60) and positively with depression symptoms (r =.39, p <.005, df = 58). These results survived controlling also for sex. Including all variables in the same analysis showed that memory and depressive symptoms were independently related to sleep problems, but neither affected the sleep-Aβ accumulation pattern. See SI for full results.

## Discussion

The present results demonstrate an age-dependent relationship between sleep problems and Aβ accumulation both globally and regionally. This fits with a view of sleep disturbances that may drive pathogenesis early in the course of neurodegeneration (38). Disturbed sleep can lead to Aβ accumulation through disruptions of sleep-dependent Aβ clearance (6), and Aβ accumulation can cause sleep problems (7), which again may reduce the brain’s ability to clear Aβ in a positive feedback loop. Several studies have reported that sleep problems are associated with accumulation of Aβ even in healthy older adults (4, 7-10, 39), but the relationships are usually relatively weak. Through direct testing of the timing of the relationship, the present results indicate that sleep problems and Aβ accumulation are more strongly related earlier than later in the normal aging process. This is interesting, as previous studies have shown that atrophy is related to sleep problems in older (> 60 years) more than in middle-aged (< 60 years) adults (40), suggesting a temporal shift in the relationship sleep and brain health for Aβ vs. atrophy.

The results were replicated by use of longitudinal information on sleep problems obtained three years prior to the present investigation. Amount of sleep problems was highly correlated between time points. Therefore, it is interesting that the participants who experienced worsening of sleep problems also had higher levels of cortical Aβ deposition, even when baseline levels of sleep problems were taken into account. Thus, both high levels of sleep problems, and increases in sleep problems regardless of the initial level, were related to Aβ deposition. Although we did not measure Aβ longitudinally, this result is in accordance with the possibility of a relationship between change in sleep problems and change in Aβ deposition. The causality could go either – or both – ways. It is less likely that there is a relationship between stable Aβ deposition and long-term increases in sleep problems, since the sleep problems would then be expected to be very severe over time. This does not fit with the observation that increased sleep problems between time points were not related to more problems at baseline. Thus, the participants with increased sleep problems may be more likely to have increased Aβ accumulation from baseline, but this needs to be tested. Recent studies have shown that even in cognitively normal older adults, Aβ deposition can increase substantially over time (41). Tracking Aβ deposition and sleep problems longitudinally over multiple time points will allow us to disentangle their age-trajectories more accurately, hopefully narrowing in on the direction of causality in human aging.

Importantly, the participants in this study are well-screened cognitively healthy older adults. Thus, we do not know who, or even if any of these will develop AD or other neurodegenerative conditions over the next decade. However, the sleep problem-related and the AD-related Aβ-accumulation showed high correspondence. This means that if sleep problems in aging are involved in Aβ-accumulation, this process to a major degree affects the same cortical regions in which AD-pathophysiology accumulates. This is in accordance with a view that sleep problems could be important in very early phases of AD (38), before clinical symptoms are detectable. The role of sleep in early AD could either be as a causal agent, as a symptom, or as a constituent in a bi-directional relationship between sleep, Aβ and neurodegeneration. However, follow-up studies over even longer time intervals are required to test whether the relationships observed in this study actually relates later development of AD or rather should be conceptualized as part of normal aging. As expected, sleep problems were also related to lower memory function and more depressive symptoms. It must be noted that one of the 21 items in BDI asks directly about sleep and another about being tired, both of which overlaps with questions of the PSQI. Still, the analyses showed that the sleep-depression and the sleep-memory relationships were independent of each other, underscoring that sleep problems are related to both cognitive function and psychiatric symptoms in aging (42).

### Gene expression – HOMER1

Sleep is a fundamental aspect of brain function, and expressions of selective genes are highly sensitive to time spent awake and time spent sleeping. Homer1a expression responds to sleep loss (16, 43), and is upregulated for both shorter (20) and longer (44) periods of sleep deprivation. The consistent activation of Homer1a suggests a role for sleep in intracellular calcium homeostasis for protecting and recovering from the neuronal activation imposed by wakefulness, and Homer1 appears to be a good marker for neuronal populations activated by sleep loss (16). Homer1’s upregulation by sleep deprivation is likely a result of such sustained neural activity, as Homer1a is transiently upregulated during increases in network activity (45). Further, the high but regionally varying expression of Homer1 in the cortex (18), and its suggested role in AD and Aβ pathophysiology (24-26), makes it a promising candidate for bridging the in vivo sleep-Aβ accumulation results, rodent studies and human brain in vitro databases.

Evidence for a role for Homer1 in Aβ processing comes from animal studies. APP+PS1 transgenic mice show reduced expression of Homer1, but normal levels in regions that do not accumulate Aβ (25). It has been suggested that inhibition of Homer 1a activity is responsible for the observed neuronal degeneration in AD by elimination of the facilitation of BK (Big Potassium) channels (26). Conversely, induction of Homer 1a can reactivate Aβ-suppressed BK channels (24). An interesting question is whether Homer1 plays a role in sleep-related Aβ accumulation in humans, and whether this again is related to AD pathophysiology. Aβ production is tightly connected to neural activity, and models are developed to explain the regional distribution of Aβ accumulation in humans as a result of regional activity variations (27). Thus, in Aβ negative healthy controls, we expected high expression of *HOMER1* in regions where high levels of AD- and sleep-related Aβ are found. This was supported by testing the overlap between the *HOMER1* gene expression map from the Allen Brain Atlas (35) and the maps of the sleep- and AD-related Aβ accumulation. These results thus may suggest a role for *HOMER1* expression in Aβ accumulation in older adults, through the critical third variable of sustained synaptic activity level, which may possibly extend to a role for Homer1 in sleep-related Aβ accumulation in preclinical AD.

### Limitations

This study has several limitations that need to be considered. Sleep problems were measured by self-report, not by polysomnography. This prevented us from testing objective measures of sleep quality, such as sleep fragmentation and amount of slow wave sleep. On the other hand, self-report is regarded as an ecologically valid way to measure sleep, since sleeping in a lab may arguably impact several aspects of sleep, and consequently be less representative of sleep patterns over time. Further, we tested *HOMER1* gene expression maps, but were not able to differentiate between different proteins encoded by this gene. The short form Homer1a is assumed to be more relevant for sleep loss than the longer forms (Homer1b and c) (16, 23). Also, although transcriptome studies may be useful in yielding a first insight into changes associated with sleep deprivation, we cannot infer from this that the specific genes are causally related to sleep, and distinguish the observed effects from changes due to secondary effects of sleep loss.

### Conclusion

Correlations between Aβ-accumulation and self-reported sleep problems have repeatedly been reported, usually ranging from weak to moderate. The present study shows that the sleep - Aβ relationship is anatomically heterogeneous and stronger in the younger-old participants than the older-old. Most importantly, the results indicate that the relationship between sleep problems and Aβ-accumulation may involve Homer**1** activity in the cortical regions that harbor Aβ in AD. This suggest a pathway through which two major AD risk factors may be causally related in older participants without clinical symptoms of AD.

## Materials and methods

### Sample

The main sample was drawn from the ongoing projects *Cognition and Plasticity through the Lifespan and Neurocognitive Plasticity* at the Center for Lifespan Changes in Brain and Cognition (LCBC), Department of Psychology, University of Oslo (46-50). All procedures were approved by the Regional Ethical Committee of Southern Norway, and written consent was obtained from all participants. Participants were mainly recruited through newspaper or internet ads. Participants were screened with a health interview, and required to be right handed, fluent Norwegian speakers, and have normal or corrected to normal vision and normal hearing. Exclusion criteria were history of injury or disease known to affect central nervous system (CNS) function, including neurological or psychiatric illness or serious head trauma, being under psychiatric treatment, use of psychoactive drugs known to affect CNS functioning, and MRI contraindications. Moreover, participants were required to score ≥26 on the Mini Mental State Examination (MMSE; (51), have a Beck Depression Inventory (BDI; (29) score ≤16, and obtain a normal IQ or above (IQ ≥ 85) on the Wechsler Abbreviated Scale of Intelligence (WASI; (52). At both time points scans were evaluated by a neuroradiologist and were required to be deemed free of significant injuries or conditions. Sample descriptives are provided in Table 1.

To calculate AD-dependent Aβ accumulation, PET scans from 20 cognitively healthy older adults (age 71.3-86.2 years, MMSE ≥ 28) and 69 patients with MCI/ AD (n = 44/ 25, 55.3-88.2 years) were obtained from the Alzheimer’s Disease Neuroimaging Initiative (ADNI) database (adni.loni.usc.edu). The ADNI was launched in 2003 as a public-private partnership, led by Principal Investigator Michael W. Weiner, MD. The primary goal of ADNI has been to test whether serial MRI, PET, other biological markers, and clinical and neuropsychological assessment can be combined to measure the progression of MCI and early AD.

### Magnetic resonance imaging acquisition and analysis

MRI data was collected using a 12-channel head coil on a 1.5 T Siemens Avanto scanner (Siemens Medical Solutions; Erlagen, Germany) at Rikshospitalet, Oslo University Hospital. The pulse sequences used included two repetitions of a 160 slices sagittal T_1_-weighted magnetization prepared rapid gradient echo (MPRAGE) sequences with the following parameters: repetition time(TR)/echo time(TE)/time to inversion(TI)/flip angle(FA)= 2400 ms/3.61 ms/1000 ms/8°, matrix = 192 × 192, field of view (FOV) = 240, voxel size = 1.25 × 1.25 × 1.20 mm, scan time 4min 42s. Cortical surfaces were reconstructed by use of FreeSurfer v. 5.3 (http://surfer.nmr.mgh.harvard.edu/) (53-55).

### Positron Emission Tomography

The participants in the main sample underwent Flutemetamol (^18^F) positron emission tomography (PET) for quantification of cortical Aβ accumulation. PET data was processed and partial voluming corrected by use of the Muller-Gartner method, registered to the individual participant’s cortical surface, and cortical surface-based smoothing (full-width, half-maximum = 15 mm) applied, shown to reduce bias and variance in PET measurements (30). The cortical PET signal at each surface was divided by the mean signal of the cerebellum cortex to obtain SUV. For some analyses, the signal in 34 cortical regions (Desikan Killian-parcellation) (31) was used. For the ADNI participants, PIB-PET images were acquired according to protocol (56, 57).

### Sleep assessment

Sleep quality was assessed using the Pittsburgh Sleep Quality Inventory (PSQI)(28) in Norwegian. PSQI is a well-validated self-rated questionnaire that assesses seven domains of sleep quality (sleep quality, latency, duration, efficiency, problems, medication and daytime tiredness) in addition to a global score over a 1-month time interval. The minimum score is 0 and maximum score is 3 for each domain, while the global score ranges from 0 to 21.

### HOMER1 expression

*HOMER1* mRNA expression levels in each of the 34 cortical regions from the Desikan-Killian atlas was extracted from the Allen Brain Atlas for the left hemisphere for six participants < 60 years (35) without know cerebral pathology (see SI), as described in detail elsewhere (36). For each statistical analysis, the median donor expression was correlated with the effect size map (gamma values) across the 34 regions. Significance was judged by each of two criteria (1) the p-value of the correlation (<.05), and (2) the position of the observed correlation among the correlations with all gene expression maps from the atlas (> 97.5 percentile).

### Statistical analyses

Surface results were tested against an empirical null distribution of maximum cluster size across 10 000 iterations using Z Monte Carlo simulations, synthesized with a cluster-forming threshold of p < 0.05 (two-sided), yielding results corrected for multiple comparisons across space.

## Acknowledgement

This work was supported by the National Health Association (to A.M.F.), Department of Psychology, University of Oslo (to K.B.W., A.M.F.), the Norwegian Research Council (to K.B.W., A.M.F.) and the project has received funding from the European Research Council’s Starting Grant and Consolidator Grant scheme under grant agreements 283634 and 725025 (to AMF) and 313440 (to KBW). The MCI/AD PET data collection and sharing was funded by the Alzheimer’s Disease Neuroimaging Initiative (ADNI) (National Institutes of Health Grant U01 AG024904) and DOD ADNI (Department of Defense award number W81XWH-12-2-0012). ADNI is funded by the National Institute on Aging, the National Institute of Biomedical Imaging and Bioengineering, and through generous contributions from the following: AbbVie, Alzheimer’s Association; Alzheimer’s Drug Discovery Foundation; Araclon Biotech; BioClinica, Inc.; Biogen; Bristol-Myers Squibb Company; CereSpir, Inc.; Cogstate; Eisai Inc.; Elan Pharmaceuticals, Inc.; Eli Lilly and Company; EuroImmun; F. Hoffmann-La Roche Ltd and its affiliated company Genentech, Inc.; Fujirebio; GE Healthcare; IXICO Ltd.; Janssen Alzheimer Immunotherapy Research & Development, LLC.; Johnson & Johnson Pharmaceutical Research & Development LLC.; Lumosity; Lundbeck; Merck & Co., Inc.; Meso Scale Diagnostics, LLC.; NeuroRx Research; Neurotrack Technologies; Novartis Pharmaceuticals Corporation; Pfizer Inc.; Piramal Imaging; Servier; Takeda Pharmaceutical Company; and Transition Therapeutics. The Canadian Institutes of Health Research is providing funds to support ADNI clinical sites in Canada. Private sector contributions are facilitated by the Foundation for the National Institutes of Health (www.fnih.org). The grantee organization is the Northern California Institute for Research and Education, and the study is coordinated by the Alzheimer’s Therapeutic Research Institute at the University of Southern California. ADNI data are disseminated by the Laboratory for Neuro Imaging at the University of Southern California.

## References

1. Hatfield CF, Herbert J, van Someren EJ, Hodges JR, & Hastings MH (2004) Disrupted daily activity/rest cycles in relation to daily cortisol rhythms of home-dwelling patients with early Alzheimer’s dementia. Brain 127(Pt 5):1061–1074.

2. Videnovic A, Lazar AS, Barker RA, & Overeem S (2014) ‘The clocks that time us’--circadian rhythms in neurodegenerative disorders. Nat Rev Neurol 10(12):683–693.

3. Prinz PN, et al. (1982) Sleep, EEG and mental function changes in senile dementia of the Alzheimer’s type. Neurobiol Aging 3(4):361–370.

4. Mander BA, et al. (2015) beta-amyloid disrupts human NREM slow waves and related hippocampus-dependent memory consolidation. Nat Neurosci 18(7):1051–1057.

5. Scullin MK & Bliwise DL (2015) Sleep, cognition, and normal aging: integrating a half century of multidisciplinary research. Perspect Psychol Sci 10(1):97–137.

6. Xie L, et al. (2013) Sleep drives metabolite clearance from the adult brain. Science 342(6156):373–377.

7. Brown BM, et al. (2016) The Relationship between Sleep Quality and Brain Amyloid Burden. Sleep 39(5):1063–1068.

8. Branger P, et al. (2016) Relationships between sleep quality and brain volume, metabolism, and amyloid deposition in late adulthood. Neurobiol Aging 41:107–114.

9. Sprecher KE, et al. (2015) Amyloid burden is associated with self-reported sleep in nondemented late middle-aged adults. Neurobiol Aging 36(9):2568–2576.

10. Spira AP, et al. (2013) Self-reported sleep and beta-amyloid deposition in community-dwelling older adults. JAMA Neurol 70(12):1537–1543.

11. Jack CR, Jr., et al. (2013) Tracking pathophysiological processes in Alzheimer’s disease: an updated hypothetical model of dynamic biomarkers. Lancet Neurol 12(2):207–216.

12. Jack CR, Jr., et al. (2010) Hypothetical model of dynamic biomarkers of the Alzheimer’s pathological cascade. Lancet Neurol 9(1):119–128.

13. Sperling RA, et al. (2011) Toward defining the preclinical stages of Alzheimer’s disease: recommendations from the National Institute on Aging-Alzheimer’s Association workgroups on diagnostic guidelines for Alzheimer’s disease. Alzheimers Dement 7(3):280–292.

14. Sperling RA, Jack CR, Jr., & Aisen PS (2011) Testing the right target and right drug at the right stage. Sci Transl Med 3(111):111cm133.

15. Pillai V, Roth T, & Drake CL (2015) The nature of stable insomnia phenotypes. Sleep 38(1):127– 138.

16. Maret S, et al. (2007) Homer1a is a core brain molecular correlate of sleep loss. Proc Natl Acad Sci U S A 104(50):20090–20095.

17. Archer SN & Oster H (2015) How sleep and wakefulness influence circadian rhythmicity: effects of insufficient and mistimed sleep on the animal and human transcriptome. J Sleep Res 24(5):476–493.

18. Szumlinski KK, Kalivas PW, & Worley PF (2006) Homer proteins: implications for neuropsychiatric disorders. Curr Opin Neurobiol 16(3):251–257.

19. Cirelli C, Faraguna U, & Tononi G (2006) Changes in brain gene expression after long-term sleep deprivation. J Neurochem 98(5):1632–1645.

20. Mackiewicz M, et al. (2007) Macromolecule biosynthesis: a key function of sleep. Physiol Genomics 31(3):441–457.

21. Thompson CL, et al. (2010) Molecular and anatomical signatures of sleep deprivation in the mouse brain. Front Neurosci 4:165.

22. Brakeman PR, et al. (1997) Homer: a protein that selectively binds metabotropic glutamate receptors. Nature 386(6622):284–288.

23. Diering GH, et al. (2017) Homer1a drives homeostatic scaling-down of excitatory synapses during sleep. Science 355(6324):511–515.

24. Luo P, Li X, Fei Z, & Poon W (2012) Scaffold protein Homer 1: implications for neurological diseases. Neurochem Int 61(5):731–738.

25. Dickey CA, et al. (2003) Selectively reduced expression of synaptic plasticity-related genes in amyloid precursor protein + presenilin-1 transgenic mice. J Neurosci 23(12):5219–5226.

26. Yamamoto K, et al. (2011) Suppression of a neocortical potassium channel activity by intracellular amyloid-beta and its rescue with Homer1a. J Neurosci 31(31):11100–11109.

27. Jagust WJ & Mormino EC (2011) Lifespan brain activity, beta-amyloid, and Alzheimer’s disease. Trends Cogn Sci 15(11):520–526.

28. Buysse DJ, Reynolds CF, 3rd, Monk TH, Berman SR, & Kupfer DJ (1989) The Pittsburgh Sleep Quality Index: a new instrument for psychiatric practice and research. Psychiatry Res 28(2):193– 213.

29. Beck AT & Steer R (1987) Beck Depression Inventory Scoring Manual (The Psychological Corporation, New York).

30. Greve DN, et al. (2014) Cortical surface-based analysis reduces bias and variance in kinetic modeling of brain PET data. Neuroimage 92:225–236.

31. Fischl B, et al. (2004) Automatically parcellating the human cerebral cortex. Cereb Cortex 14(1):11–22.

32. Beck AT & Steer RA (1984) Internal consistencies of the original and revised Beck Depression Inventory. J Clin Psychol 40(6):1365–1367.

33. Vorona RD, et al. (2005) Overweight and obese patients in a primary care population report less sleep than patients with a normal body mass index. Arch Intern Med 165(1):25–30.

34. Ling CH, et al. (2011) Accuracy of direct segmental multi-frequency bioimpedance analysis in the assessment of total body and segmental body composition in middle-aged adult population. Clin Nutr 30(5):610–615.

35. Hawrylycz MJ, et al. (2012) An anatomically comprehensive atlas of the adult human brain transcriptome. Nature 489(7416):391–399.

36. French L & Paus T (2015) A FreeSurfer view of the cortical transcriptome generated from the Allen Human Brain Atlas. Front Neurosci 9:323.

37. Poulton RG & Moffitt TE (1995) The Rey-Osterreith Complex Figure Test: norms for young adolescents and an examination of validity. Arch Clin Neuropsychol 10(1):47–56.

38. Musiek ES & Holtzman DM (2016) Mechanisms linking circadian clocks, sleep, and neurodegeneration. Science 354(6315):1004–1008.

39. Ju YE, et al. (2013) Sleep quality and preclinical Alzheimer disease. JAMA Neurol 70(5):587–593.

40. Sexton CE, Storsve AB, Walhovd KB, Johansen-Berg H, & Fjell AM (2014) Poor sleep quality is associated with increased cortical atrophy in community-dwelling adults. Neurology 83(11):967– 973.

41. Resnick SM, et al. (2015) Changes in Abeta biomarkers and associations with APOE genotype in 2 longitudinal cohorts. Neurobiol Aging 36(8):2333–2339.

42. Mander BA, Winer JR, & Walker MP (2017) Sleep and Human Aging. Neuron 94(1):19-36.

43. Wang H, Liu Y, Briesemann M, & Yan J (2010) Computational analysis of gene regulation in animal sleep deprivation. Physiol Genomics 42(3):427–436.

44. Conti B, et al. (2007) Region-specific transcriptional changes following the three antidepressant treatments electro convulsive therapy, sleep deprivation and fluoxetine. Mol Psychiatry 12(2):167–189.

45. Hu JH, et al. (2010) Homeostatic scaling requires group I mGluR activation mediated by Homer1a. Neuron 68(6):1128–1142.

46. Westlye LT, Grydeland H, Walhovd KB, & Fjell AM (2010) Associations between Regional Cortical Thickness and Attentional Networks as Measured by the Attention Network Test. Cerebral cortex 21(2):345–356.

47. Westlye LT, et al. (2010) Differentiating maturational and aging-related changes of the cerebral cortex by use of thickness and signal intensity. Neuroimage 52(1):172–185.

48. Storsve AB, et al. (2014) Differential longitudinal changes in cortical thickness, surface area and volume across the adult life span: regions of accelerating and decelerating change. J Neurosci 34(25):8488–8498.

49. Walhovd KB, Storsve AB, Westlye LT, Drevon CA, & Fjell AM (2014) Blood markers of fatty acids and vitamin D, cardiovascular measures, body mass index, and physical activity relate to longitudinal cortical thinning in normal aging. Neurobiol Aging 35(5):1055–1064.

50. de Lange AG, et al. (2016) White matter integrity as a marker for cognitive plasticity in aging. Neurobiol Aging 47:74–82.

51. Folstein MF, Folstein SE, & McHugh PR (1975) ‚ÄúMini-mental state‚Äù: A practical method for grading the cognitive state of patients for the clinician. Journal of Psychiatric Research 12(3):189–198.

52. Wechsler D (1999) Wechsler abbreviated scale of intelligence (The Psychological Corporation, San Antonio, TX).

53. Dale AM, Fischl B, & Sereno MI (1999) Cortical surface-based analysis. I. Segmentation and surface reconstruction. NeuroImage 9:179–194.

54. Fischl B, Sereno MI, & Dale AM (1999) Cortical surface-based analysis. II: Inflation, flattening, and a surface-based coordinate system. NeuroImage 9:195–207.

55. Fischl B & Dale AM (2000) Measuring the thickness of the human cerebral cortex from magnetic resonance images. Proc. Natl. Acad. Sci. U S A 97:11050–11055.

56. Jack CR, et al. (2008) The Alzheimer’s Disease Neuroimaging Initiative (ADNI): MRI Methods. Journal of magnetic resonance imaging: JMRI 27(4):685–691.

57. Jagust WJ, et al. (2010) The ADNI PET Core. Alzheimer’s & dementia: the journal of the Alzheimer’s Association 6(3):221–229.

